# Decoding strawberry volatile cultivar diversity through comparative transcriptome analysis

**DOI:** 10.1101/2024.03.14.584999

**Authors:** Kondylia Passa, Evangelos Tsormpatsidis, Ioannis Ganopoulos, Christos Bazakos, Vasileios Papasotiropoulos

## Abstract

This study presents a comparative transcriptomic analysis of three commercial strawberry cultivars: ‘Rociera’, ‘Calderon’, and ‘Victory’, aimed at uncovering the molecular basis of their distinct flavor and aroma profiles. Through RNA sequencing, we analyzed the transcriptomic landscape of these varieties, uncovering a notable array of differentially expressed genes (DEGs) between them. Specifically, ‘Rociera’ showed a higher count of DEGs compared to ‘Victory’, suggesting significant differences in gene expression related to flavor and aroma between those cultivars. Our further investigations revealed pivotal metabolic pathways— such as those involving furanones, amino acids, fatty acids, and terpenoids—participating in generating volatile organic compounds in strawberries. These pathways exhibited variety-specific expression patterns, underlining the genetic determinants of each variety’s unique sensory characteristics. Gene Ontology (GO) enrichment analysis underscored important pathways like phototropism, sugar-mediated signaling, and terpenoid biosynthesis, highlighting the genetic intricacy that influences strawberry flavor and aroma. Additionally, our study identified 31,677 single nucleotide polymorphisms (SNPs) with a significant impact on genes associated with volatile compound biosynthesis. Notably, SNPs linked to key enzymes such as lipoxygenase 6, omega-6 fatty acid desaturase (FaFAD1), and phenylalanine ammonia-lyase 1 (PAL) were discovered, shedding light on the genetic variations that underlie flavor differences. Therefore, this research advances our comprehension of the genetic elements that impact strawberry fruit flavor and aroma, offering invaluable insights for molecular breeding endeavors focused on enhancing strawberry flavor.

## Introduction

Strawberries, cherished worldwide for their unique flavor and aroma, are cultivated globally not only for these traits, but also for their rich nutritional profile, which includes a variety of health-beneficial compounds [1,2]. The commercial strawberry (*Fragaria* × *ananassa* (Duchesne ex Weston) Duchesne ex Rozier), an octoploid species within the *Rosaceae* family, originated from a cross between North American *Fragaria virginiana* and South American *Fragaria chiloensis* [3]. As one of the most economically significant berries, strawberries achieved a global production of 9.57 million metric tons in 2022 [4].

The distinctive flavor and aroma of strawberries, attributed to sugars and a complex mixture of volatile organic compounds (VOCs), play a crucial role in consumer preference and the overall appeal of this fruit [5]. Key components of strawberry aroma include esters, lactones, acids, furanones, terpenes, and sulfur compounds, with specific metabolites such as ethyl butanoate, methyl butanoate, mesifurane (DMMF), furaneol (DMHF), linalool, nerolidol, γ-decalactone, butanoic acid, and hexanoic acid significantly contributing to the organoleptic profile of strawberries [6–10]. The biosynthesis of these secondary metabolites is regulated through various biochemical pathways, including amino acid metabolism, fatty acid metabolism, and terpenoid metabolism [11,12].

Over recent decades, numerous genes and genetic loci associated with strawberry volatiles have been identified, such as the *SAAT* gene encoding alcohol acyltransferase (AAT) for volatile ester synthesis [13], *FaQR* involved in furaneol biosynthesis, and *FaOMT* controlling mesifurane content [14–16]. The fatty acid desaturase gene *FaFAD1* influences γ-decalactone production in ripening fruits [17], while eugenol synthases (*FaEGS1* and *FaEGS2*) are pivotal for eugenol production in strawberry achenes and receptacles [18]. The transcription factor FaEOBII and *FaMYB10* play significant roles in regulating the phenylpropanoid pathway and the expression of key genes in eugenol biosynthesis [19,20]. *F. ananassa* nerolidol synthase1 (*FaNES1*) is responsible for synthesizing the monoterpenes and sesquiterpenes linalool and nerolidol [13]. Research has also focused on factors influencing strawberry fruit’s volatile profile, including cultivation practices, environmental conditions, fruit maturity stage, postharvest treatments, and genetic/cultivar differences [20–22]. Transcriptome analysis and gene expression studies have provided insights into the volatile profile variations between commercial strawberry cultivars, wild accessions, or their somaclonal mutants at different ripening stages [24,25]. RNA sequencing has emerged as an effective tool for unraveling the regulatory mechanisms involved in fruit development and ripening, offering deeper insights into the genes central to these biochemical pathways [26–28]. Chambers *et al*. (2014) combined transcriptomic and analytical chemistry techniques to identify *FaFAD1*, a gene likely influencing a key flavor volatile in strawberries [29].

This study aims to delve into the genetic and molecular basis underlying the flavor and aroma variations in three commercial strawberry cultivars: ‘Rociera’, ‘Calderon’, and ‘Victory’. By conducting transcriptomic analyses, we pinpointed differentially expressed genes (DEGs) and genetic variations that contribute to these cultivars’ sensory traits, intending to guide molecular breeding strategies toward developing strawberry varieties with enhanced flavor and aroma profiles tailored to consumer preferences.

## Materials and Methods

### Plant Material

The present study was conducted during the 2022/2023 growing season at Berryplasma World Ltd., Varda Ilias (38°01’50.09” N 21°21’54.22” E), Region of Western Greece. Three different commercial cultivars, ‘Rociera’, ‘Calderon’ and ‘Victory’, were cultivated at the same plantation under the same conditions.

‘Rociera’ was developed by Fresas Nuevos Materiales S.A. (FNM) (https://www.fresasnm.com/) in Spain, ‘Calderon’ by Masiá Ciscar S.A. (https://www.masiaciscar.es/) in Spain and ‘Victory’ by Berry Genetics® (https://www.berrygenetics.com/) in Interlaken, Santa Cruz, CA, USA. The experiment was carried out in commercial tunnels, each measuring 8.2 × 3.5 × 6.8 m (W × H × L). Mother plants of each variety were planted in 10L pots in summer 2022 and grown until they produced runner tips. All runner tips from each variety were harvested in July 2022 and plugged into trays. All misted tips were grown under the same conditions until planting in plastic tunnels. Strawberry plants were irrigated with a standard commercial nutrient solution applied through a drip irrigation system via a Dosatron (Dosatron International, Bordeaux, France) set at an electrical conductivity of 1.85 mS (N: 120 ppm, P: 50 ppm, K: 180 ppm, Mg: 30 ppm, Ca: 100 ppm, Fe: 5 ppm, Mn: 0.15 ppm, Zn: 0.15 ppm, B: 0.2 ppm, Cu: 0.02 ppm, Mo: 0.015 ppm, pH: 6.00). Plug plants of the three cultivars were planted in 1 M coir growbags (Dutch Plantin, India) and were placed in raised beds at a density of 5 plants per growbag and 60,000 plants per ha in total.

Fruits from three different plants of each cultivar were collected and the achenes were removed from the receptacle. Each replicate sample, consisting of 50mg of fresh tissue from each individual plant, was placed in separate Eppendorf tubes. The tubes with the tissues, as well as the rest of the strawberry fruits collected, were immersed in liquid nitrogen, and were stored at −80°C.

### Total RNA extraction, library construction and RNA-Seq analysis

Total RNA was isolated by using the RNeasy® Mini Kit from Qiagen (Valencia, CA, USA), according, to the manufacturer’s protocol. The integrity and quality of the extracted RNA were accessed via Agilent 2100 Bioanalyzer (Agilent, CA, USA). Subsequently, nine Illumina-Truseq stranded libraries were prepared and sequenced by Novogene Europe (https://www.novogene.com/eu-en/) using an Illumina Novaseq 6000 sequencer. The raw sequencing data were processed to remove adaptor sequences, reads containing over 5% unknown bases (N), and reads of low quality. The processed sequences, termed clean reads, were preserved in the FASTQ format. Subsequent mapping of these short sequencing reads to the *Fragaria x ananassa* ‘Camarosa’ v1.0 reference genome [30] was performed using HISAT 2.2.1 [31] with default parameters. The resulting data were sorted, and the mapped reads were quantified using HTseq-count [32], based on the updated annotation *Fragaria x ananassa* ‘Camarosa’ Genome v1.0.a2 [33]. Expression levels were calculated as transcripts per million (TPM) [34], and the identification of differentially expressed genes among the genotypes was achieved using the edgeR package [35], with raw counts as input. This TPM-normalized data was then analyzed through Principal Component Analysis (PCA) and Hierarchical Clustering (HC). The Illumina RNA-Seq reads are available from NCBI (project PRJNA917223). Genes were identified as up-regulated if they had a p-value adjusted to ≤0.01 and a positive log2 fold change, while those with a negative log2 fold change were considered as down-regulated.

### Quantitative Real-Time PCR (qRT-PCR)

Extraction of mRNA was conducted in three biological replicates by using the RNeasy® Mini Kit from Qiagen (Valencia, CA, USA). To obtain cDNA, 500 ng RNA was reverse transcribed using the iScriptTM cDna Synthesis kit (Biorad Laboratories, Inc.) and a PCR (C1000 TouchTM Thermal Cycler, Biorad Laboratories, Inc.). First-strand cDNA synthesis of 500 ng of RNA in a final volume of 20 μL was performed using iScript cDNA synthesis kit (Bio-Rad Laboratories, Hercules, CA, USA), according to the manufacturer’s protocol. The synthesized cDNA was used as the template for qRT-PCR reactions in a total volume of 15 μL, consisting of 7 μL Sso Advanced Universal SYBR Green Supermix (Bio-Rad Laboratories, Hercules, CA, USA), 0.28 μL of each primer (10 μM), 6.94 μL of water, and 0.5 μL of cDNA on a CFX 96™ Real-Time PCR Detection System (Bio-Rad Laboratories, Hercules, CA, USA), using the corresponding specific primers [26,36,37] for the analyzed genes (*SAAT*, *FaQR*, *FaADH*). The reaction was performed with an initial denaturation step at 95 °C for 5 min, followed by 39 cycles at 95 °C for 30 s, primer annealing temperature 58 °C for 30 s and extension temperature 72 °C for 1 min, with a plate read between each cycle. A melting curve analysis was conducted between 70 and 90 °C with a read every 0.5 °C held for 2 s between each read, to verify the specificity of primer amplification, based on the presence of a single and sharp peak. Negative controls were included in all amplification reactions to check for potential reagent contamination. Data were analyzed using the ΔΔCt method and were expressed as −ΔΔCt.

### Single nucleotide polymorphism (SNP) calling and annotation

The assessment of paired-end read quality was performed using FastQC (version 0.11.9) and MultiQC (version 1.9). Adaptors were trimmed and reads were filtered to remove those of low quality (Q > 28) and with unknown nucleotides (N) using Trim Galore. Reads that met the specified quality standards (average quality ≥ Q20, length ≥ 18 bp) were then mapped to the *Fragaria x ananassa ‘* Camarosa’ v1.0 reference genome [30] utilizing Hisat2. Picard (version 2.22.8) was employed to identify PCR duplicates, and samtools (version 1.11) [38] was used to eliminate reads mapping to multiple locations. The SplitNCigarReads tool was utilized to split reads containing N into multiple supplementary alignments and hard clips mismatching overhangs. Variant calling was performed with the Genome Analysis ToolKit (GATK), using a suite of tools including HaplotypeCaller, CombineGVCFs, GenotypeGVCFs, Select Variants, and Variant Filtration, to identify and categorize small variants as either Single Nucleotide Polymorphisms (SNPs) or insertions/deletions (InDels). Variants specific to individual samples were isolated with bcftools (version 1.11), and SnpEff (version 4.3) [39] was applied for variant annotation.

## Results

### RNA-Seq analysis of three commercial strawberry cultivars

To understand the flavor and aroma profile differences among the three cultivars, we performed a transcriptomic analysis (RNA-Seq) on the flesh tissues of the commercial strawberry cultivars ‘Rociera’, ‘Calderon’, and ‘Victory’. After removing low-quality reads, we obtained an average of 33,638,614 clean reads for ‘Rociera’, 32,738,587 for ‘Calderon’ and 31,246,491 for ‘Victory’. These reads were mapped to the *Fragaria x ananassa* reference genome, achieving an average mapping rate of 81.9% (Table 1).

**Table 1.** Number of reads and aligned sequenced from RNA-Seq of the three strawberry cultivars (‘Rociera’, ‘Calderon’ and ‘Victory’). F_ROC means ‘Rociera’ strawberry fruit, F_CAL means ‘Calderon’ strawberry fruit, F_VIC means ‘Victory’ strawberry fruit, _1, _2, _3 corresponds to the biological replicate number.

| Library | Raw reads | Clean reads | Clean reads aligned to transcriptome assembly (%) |
| --- | --- | --- | --- |
| F_ROC_1 | 34463917 | 34147901 | 81.50 |
| F_ROC_2 | 34526199 | 33997287 | 81.18 |
| F_ROC_3 | 33124111 | 32770655 | 81.60 |
| F_CAL_1 | 33046778 | 32716786 | 81.64 |
| F_CAL_2 | 34053556 | 33533360 | 82.10 |
| F_CAL_3 | 32378318 | 31965614 | 82.05 |
| F_VIC_1 | 30472046 | 30025754 | 82.31 |
| F_VIC_2 | 30765651 | 30433951 | 82.60 |
| F_VIC_3 | 33764385 | 33279768 | 82.32 |

| **Library** | **Raw reads** | **Clean reads** | **Clean reads aligned to transcriptome assembly (%)** |
| --- | --- | --- | --- |
| F_ROC_1 | 34463917 | 34147901 | 81.50 |
| F_ROC_2 | 34526199 | 33997287 | 81.18 |
| F_ROC_3 | 33124111 | 32770655 | 81.60 |
| F_CAL_1 | 33046778 | 32716786 | 81.64 |
| F_CAL_2 | 34053556 | 33533360 | 82.10 |
| F_CAL_3 | 32378318 | 31965614 | 82.05 |
| F_VIC_1 | 30472046 | 30025754 | 82.31 |
| F_VIC_2 | 30765651 | 30433951 | 82.60 |
| F_VIC_3 | 33764385 | 33279768 | 82.32 |

Hierarchical clustering (HC) and PCA demonstrated a distinct separation among the three cultivars (Figure 1). ‘Rociera’ vs. ‘Victory’ comparison showed the highest number of Differentially Expressed Genes (DEGs), while ‘Victory’ vs. ‘Calderon’ exhibited the fewest. In total, 4538 and 4107 genes were up- and down-regulated, respectively, in ‘Rociera’ compared to ‘Victory’; 3483 and 3479 in ‘Rociera’ compared to ‘Calderon’; and 2776 and 3139 in ‘Victory’ compared to ‘Calderon’.

**Figure 1.**
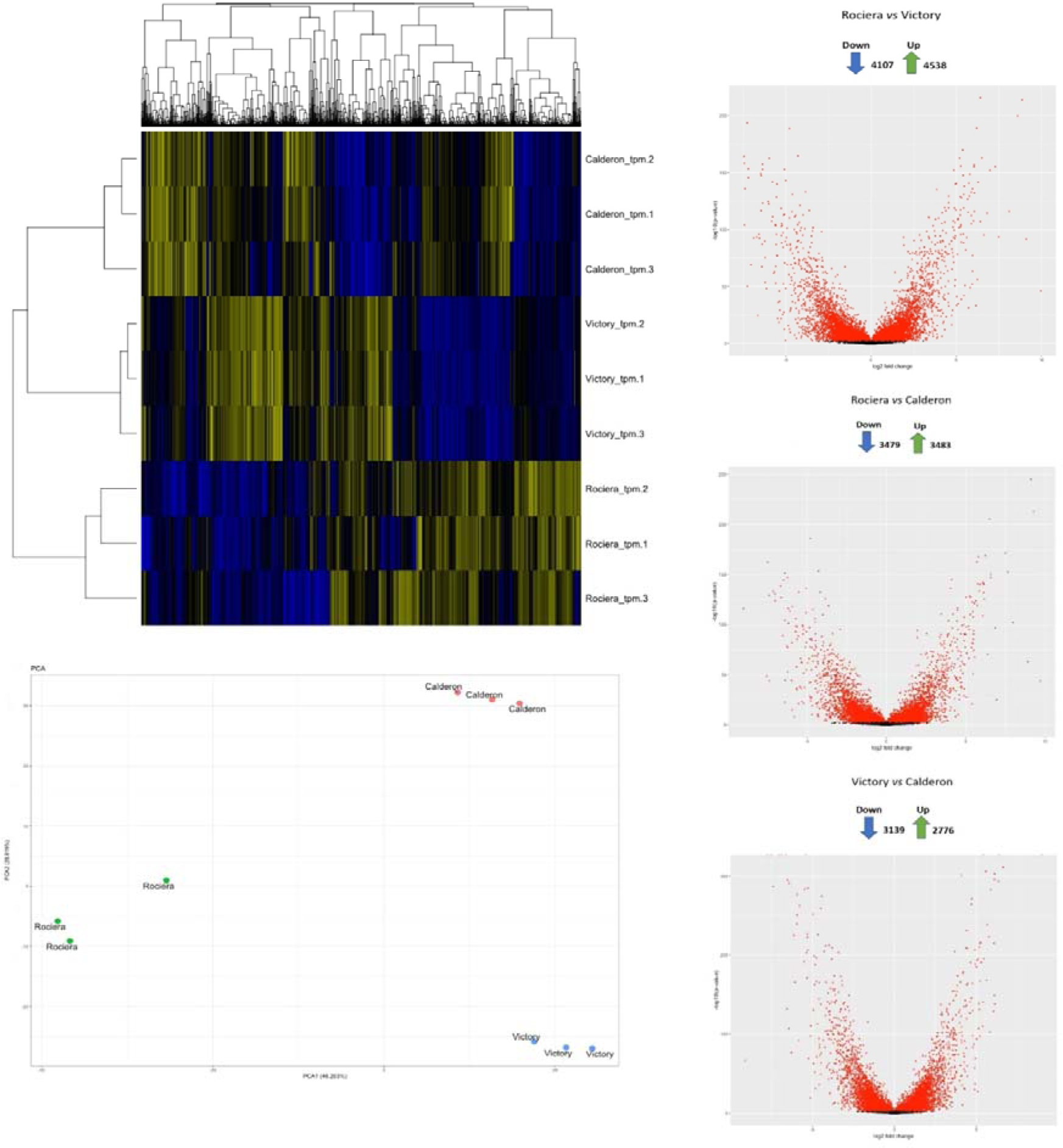
A global overview of the gene expression differences in strawberry fruits in the three commercial cultivars studied. (a) Hierarchical clustering (Euclidean distance metric/spearman correlation method) (b) Principal Component Analysis (PCA) and (c) Volcano plot depict clustering of differentially expressed (upregulated and down-regulated) genes in ‘Rociera’ (vs ‘Victory’), Rociera (vs Calderon) and Victory (vs Calderon).

To pinpoint the DEGs among the fruit samples, comparisons were made between ‘Rociera’ vs. ‘Victory’ and ‘Rociera’ vs. ‘Calderon’. In ‘Rociera’, 6,070 upregulated DEGs were identified, with 1951 common to both comparisons (Table S1; Figure 2). Venn diagrams illustrated the upregulated and downregulated DEGs in both comparisons (Figure 2), suggesting these DEGs could explain the unique aroma and flavor traits of ‘Rociera’ relative to ‘Victory’ and ‘Calderon’.

**Figure 2.**
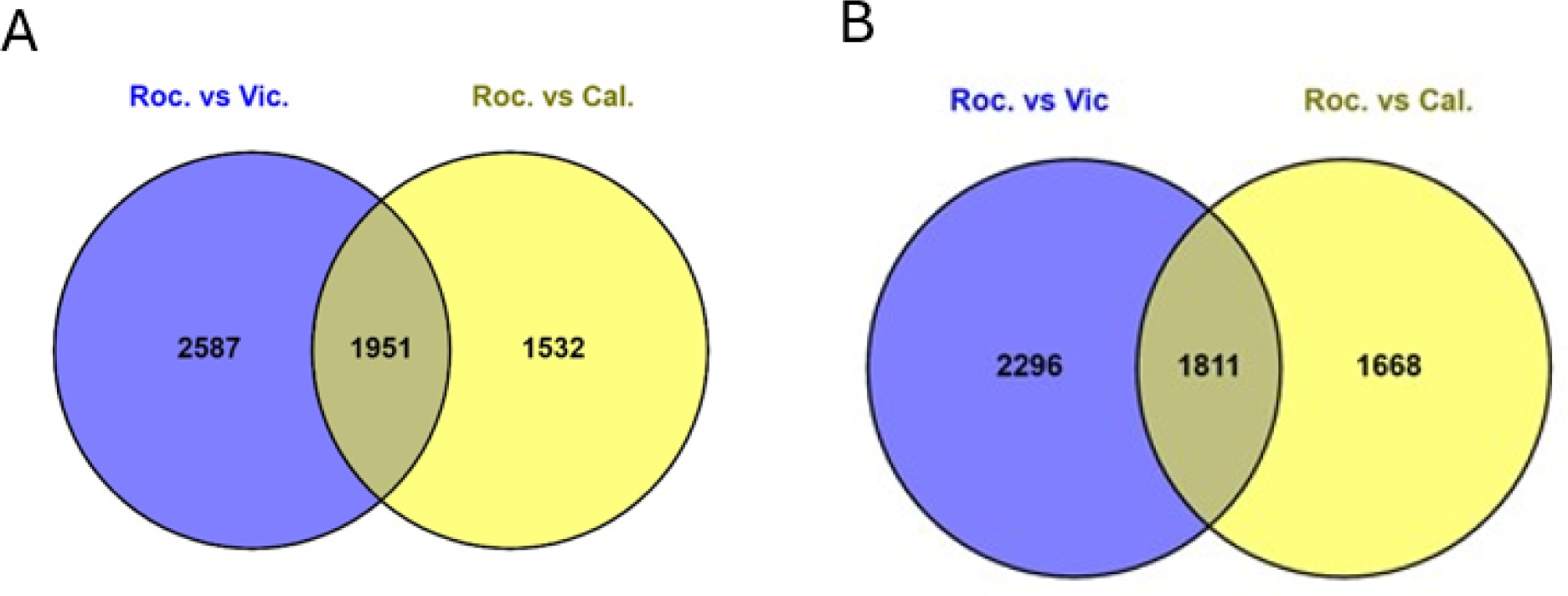
Venn diagram analysis of differentially expressed genes upregulated (A) and downregulated (B) in Rociera.

GO enrichment analysis functionally categorized the significant DEGs, identifying the 17 most enriched GO pathways among the upregulated DEGs in ‘Rociera’ vs. ‘Victory’ and ‘Rociera’ vs. ‘Calderon’ comparisons (Figure 3). The most enriched GO terms in biological process (BP) were phototropism, sugar mediated signaling pathway and carbohydrate mediated signaling and the most significant molecular functions (MF) included ATP-dependent activity and cation binding. Significant GO terms in BP enrichment were also: response to carbohydrate, diterpenoid and terpenoid biosynthetic process. Our findings indicated that genotypic differences impact the terpenoid biosynthetic pathway, influencing strawberry fruit flavor/aroma, therefore, we focused on gene groups implicated in pathways for the synthesis of strawberry volatile organic compounds. Specifically, we studied the expression of gene groups involved in amino acid, fatty acid, terpenoid and furanones pathway.

**Figure 3.**
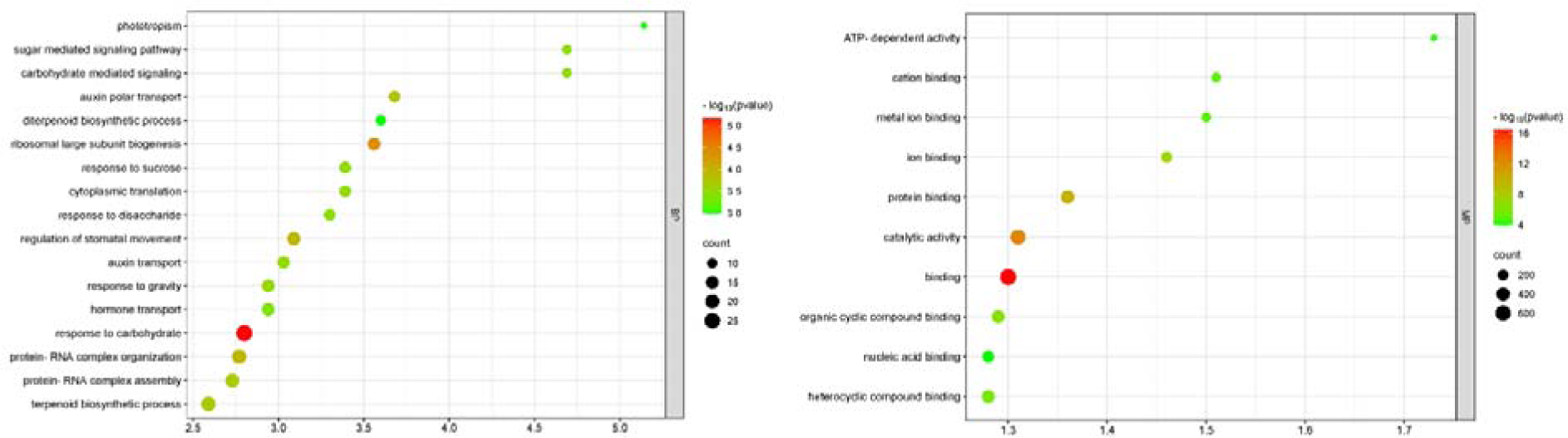
GO enrichment analysis of upregulated differentially expressed genes in Rociera.

### Furanones Pathway

The biosynthesis of furanones, including furaneol and mesifurane, originates from carbohydrate metabolism [40,41]. Research indicates that both furaneol and mesifurane (DMMF) naturally derive from D-fructose-1,6-diphosphate [42]. The *Fragaria x ananassa* gene for quinone oxireductase (*FaQR*) plays a crucial role in the biosynthetic pathway leading to furaneol (DMHF), with 4-Hydroxy-5-methyl-2-methylene-3(2H)-furanone (HMMF) identified as a precursor to HDMF and a natural substrate for *FaQR*. Recent findings suggest that the transcription factor complex, including *FaERF*#9 and *FaMYB98*, activates the *FaQR* promoter, thereby enhancing furaneol biosynthesis in strawberries [15,16]. The enzymatic methylation of furaneol (DMHF) by O-methyltransferase (FaOMT) results in the formation of mesifurane (DMMF) [43]. *FaOMT* has been pinpointed as the gene responsible for the natural variation in DMMF content within strawberry fruits [14] (Figure 4A).

**Figure 4:**
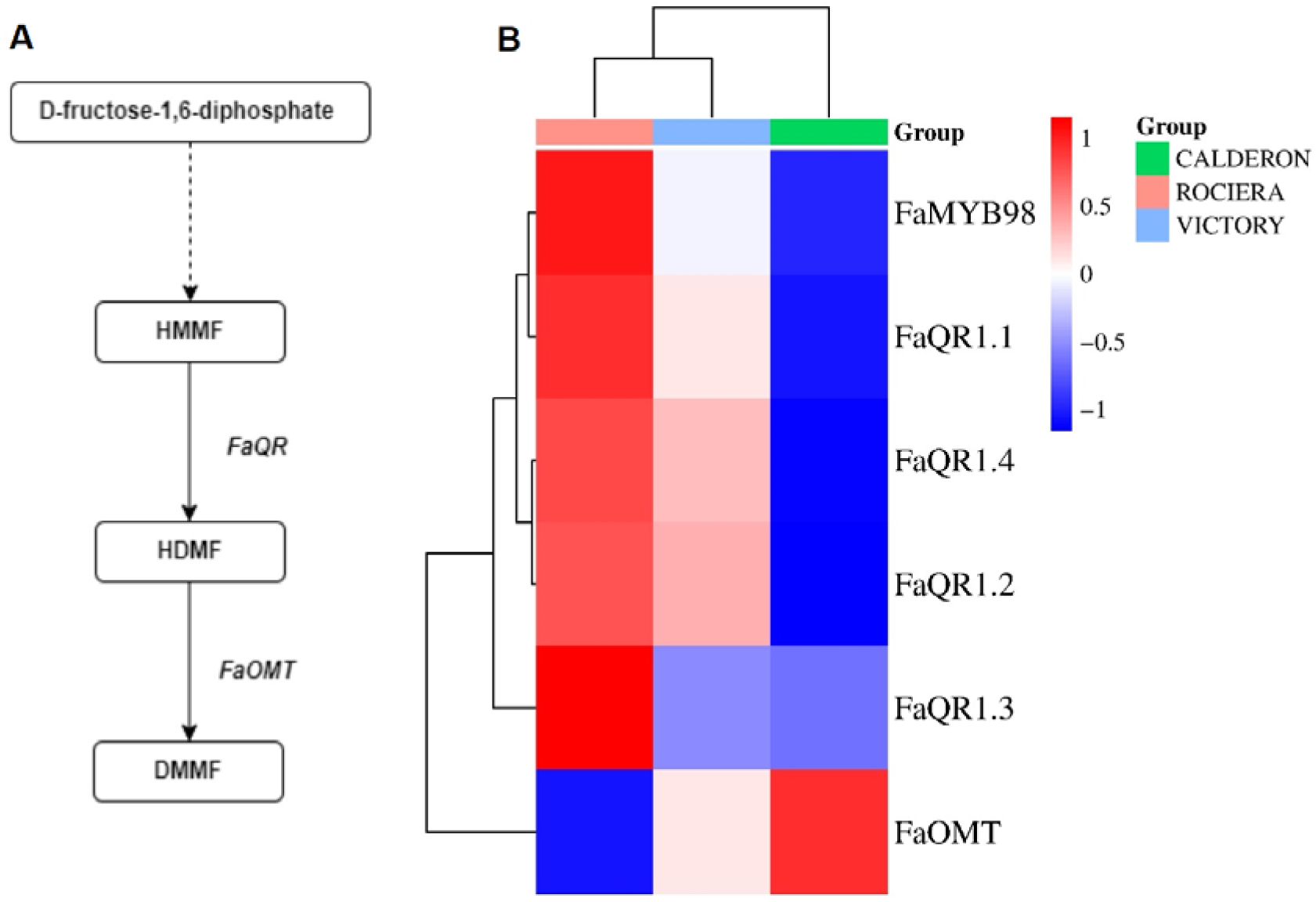
Furanones pathway analysis. (A) Schematic diagram of furanones pathways. (B) Transcript level of biosynthetic genes related to furanones pathway in the fruit of ‘Rociera’ ‘Victory’ and ‘Calderon’ varieties. Darker red indicates a higher expression level, whereas darker blue indicates a lower expression level.

Significant variation was observed in the expression levels of the quinone oxidoreductase gene *FaQR*, which is instrumental in synthesizing furaneol (2,5-dimethyl-4-hydroxy-3(2H)-furanone, DMHF), among the three examined varieties. ‘Rociera’ showed an upregulation of *FaQR* expression, in contrast to ‘Calderon’. Additionally, the transcription factor *FaMYB98* was found to be upregulated in ‘Rociera’. Inversely, *FaOMT* expression was reduced in ‘Rociera’ while being elevated in ‘Calderon’ (Figure 4B).

### Amino acid and fatty acid pathways

Volatile esters are synthesized via fatty acid and amino acid pathways, with aldehydes formed by LOX and HPL activity in the former, and PDC activity in the latter [25,44]. Alcohol acyltransferase (AAT) enzyme is crucial for esterification (Figure 5A). *FaAAT* was specifically upregulated in ‘Rociera’, hinting at its stronger aroma profile (Figure 5B).

**Figure 5:**
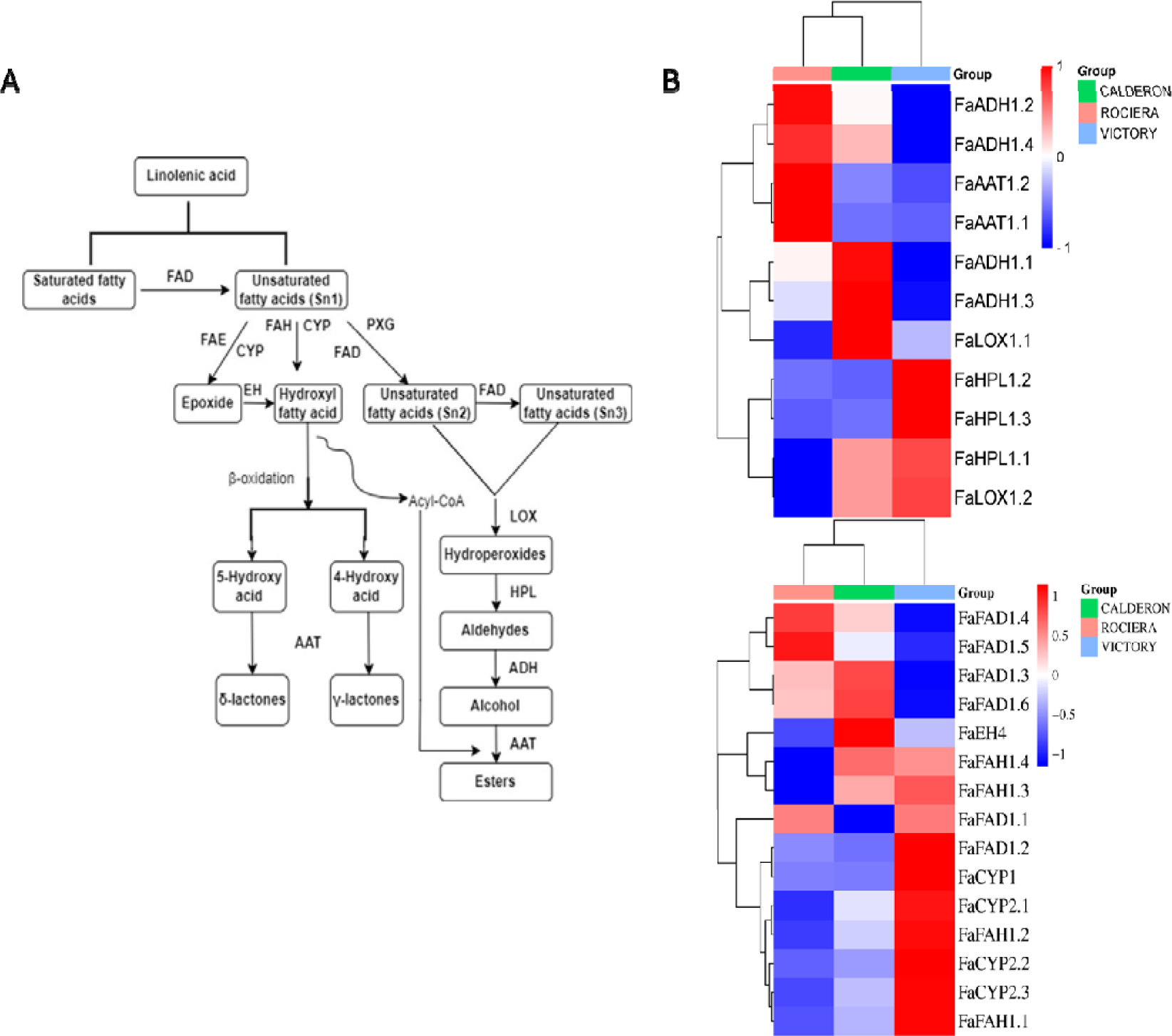
Fatty acid pathway analysis. (**A**) Schematic diagram of the fatty acid pathway. Multiple enzymatic reactions are illustrated by stacked arrows. Abbreviations: ACX: acyl-CoA oxidases; FAD: fatty acid desaturase; FAE: fatty acid elongase; HPL: hydroperoxide lyase; CYP: cytochrome P450; FAH: fatty acid hydroxylase; PXG: peroxygenase; EH: epoxide hydrolase; LOX: lipoxygenase; ADH: alcohol dehydrogenases; AAT: alcohol acyl-transferases; (**B**) Transcript level of biosynthetic genes related to fatty acid pathway in the fruit of ‘Rociera’ ‘Victory’ and ‘Calderon’ varieties. Darker red indicates a higher expression level, whereas darker blue indicates a lower expression level.

Lactone synthesis originates from the β-oxidation of unsaturated fatty acids and involves a series of enzymes known as desaturases (FADs) [17,45] (Figure 5A). Specifically, the fatty acid desaturase (FaFAD1) has been identified as a key enzyme positively associated with the synthesis of γ-decalactone during the ripening of strawberries [17,29,46]. For eugenol biosynthesis, L-phenylalanine serves as the starting molecule, leading to eugenol through several enzymatic reactions facilitated by enzymes such as phenylalanine ammonia-lyase (PAL), cinnamate-4-hydroxylase (C4H), p-coumarate 3-hydroxylase (C3H), caffeic acid 3-O-methyltransferase (COMT), 4-coumarate-CoA ligase (4CL), cinnamoyl-CoA reductase (CCR), cinnamyl alcohol dehydrogenase (CAD), and coniferyl alcohol acetyltransferase (CFAT), culminating with eugenol synthase (EGS) in the amino acid pathway [47]. The transcription factor *FaMYB10*, pivotal in the flavonoid/phenylpropanoid pathway, influences the expression of *FaEOBII*, which in turn activates crucial genes for eugenol production [19] (Figure 6A).

**Figure 6:**
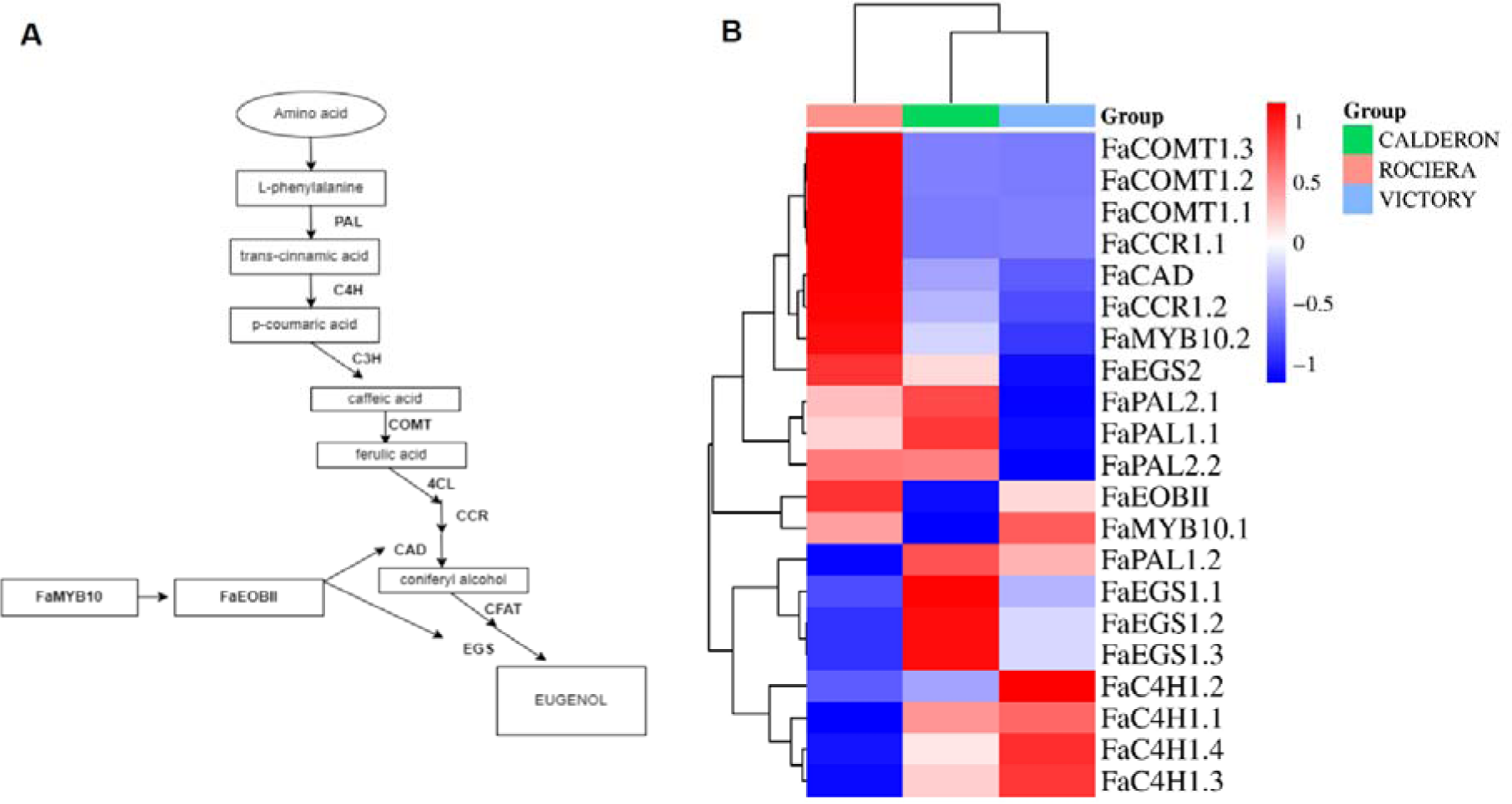
Amino acid pathway analysis. (**A**) Schematic diagram of the amino acid pathway. Multiple enzymatic reactions are illustrated by stacked arrows. Abbreviations: PAL, Phenylalanine ammonia lyase; C4H, Cinnamate 4-hydroxylase; C3H, p-coumarate 3-hydroxylase; COMT, caffeic O-methyltransferase; 4CL, 4-coumarate: CoA ligase; CCR, cinnamoyl CoA reductase; CAD, cinnamyl alcohol dehydrogenase; CFAT, Coniferyl alcohol acetyltransferase. (**B**) Transcript level of biosynthetic genes related to amino acid pathway in the fruit of ‘Rociera’ ‘Victory’ and ‘Calderon’ varieties. Darker red indicates a higher expression level, whereas darker blue indicates a lower expression level.

The lipoxygenase gene *FaLOX* is downregulated in ‘Rociera’ while showing upregulation in ‘Calderon’. Conversely, hydroperoxide lyase (*FaHPL*) was upregulated in ‘Victory’. The alcohol dehydrogenase genes *FaADH1.2* and *FaADH1.4* were upregulated in ‘Rociera’, whereas *FaADH1.1* and *FaADH1.3* in ‘Calderon’. Notably, ‘Rociera’ uniquely displayed upregulation of the ester production gene *FaAAT* (Figure 5B). In the context of lactone biosynthesis, *FaFAH* showed upregulation in ‘Victory’ but downregulation in ‘Rociera’, with a similar expression pattern observed in *FaCYP* transcript levels. *FaEH4* was upregulated in ‘Calderon’. Among the desaturases, *FaFAD1*.*3*, *FaFAD1.4*, *FaFAD1.5*, and *FaFAD1.6* were downregulated in ‘Victory’ but upregulated in both ‘Rociera’ and ‘Calderon’, while *FaFAD1.1* and *FaFAD1.2* were upregulated in ‘Victory’ (Figure 5Β).

In relation to eugenol biosynthesis, *FaPAL* was upregulated in ‘Calderon’ and ‘Rociera’ but downregulated in ‘Victory’. *FaC4H* saw increased expression in ‘Victory’ but was reduced in ‘Rociera’. The caffeic O-methyltransferase genes were exclusively upregulated in ‘Rociera’, displaying a downregulated expression pattern in both ‘Victory’ and ‘Calderon’. Both *FaCCR* and *FaCAD* were upregulated in ‘Rociera’, whereas these genes were downregulated in ‘Victory’ and ‘Calderon’. Interestingly, *FaEGS1* and *FaEGS2* showed contrasting expression patterns, with *FaEGS1* downregulated and *FaEGS2* upregulated in ‘Victory’ and ‘Calderon’. In ‘Rociera’, *FaEGS1* was downregulated, yet *FaEGS2*, along with transcription factors *FaMYB10* and *FaEOBII*, exhibited high expression levels (Figure 6B).

### Terpenoid Pathway

Dimethylallyl diphosphate (DMAPP), produced through the 2-C-methyl-D-erythritol 4-phosphate (MEP) pathway, and isopentenyl pyrophosphate (IPP), synthesized via both the MEP and mevalonate (MVA) pathways, serve as the foundational precursors for terpenoids [48]. The enzyme 1-deoxy-D-xylulose 5-phosphate synthase (DXS) facilitates the condensation of pyruvate with D-glyceraldehyde 3-phosphate (G-3P). The reduction of 1-deoxy-D-xylulose 5-phosphate (DXP) is driven by DXP reductoisomerase (DXR). This pathway includes several enzymes, such as 2-C-methyl-D-erythritol 4-phosphate cytidylyltransferase (CMS), 4- (cytidine 51-diphospho)-2-C-methyl-D-erythritol kinase (CMK), 2-C-methyl-D-erythritol 2, 4-cyclodiphosphate synthase (MCS), 4-hydroxy-3-methylbut-2-en-1-yl diphosphate synthase (HDS), and HMBPP (4-hydroxy-3-methylbut-2-enyldiphosphate) reductase (HDR). Terpene synthases (TPSs) conclude the terpenoid synthesis process. Specifically, nerolidol synthase (*FaNES1*), which participates in synthesizing nerolidol and linalool, is predominantly active in the strawberry receptacle during fruit ripening [13] (Figure 7A). Among the cultivars, ‘Calderon’ showed exclusive upregulation of the *FaDXS* gene, whereas the same was also observed in ‘Rociera’ for *FaDXR*. On the other hand, ‘Victory’ exhibited the highest expression levels for *FaCMS*; both *FaHDS* and *FaHDR* were similarly downregulated in ‘Victory’, while in ‘Rociera’, expression of *FaHDR* and *FaTPS* was increased. *FaNES1* was upregulated in both ‘Rociera’ and ‘Victory’ while it was downregulated in ‘Calderon’ (Figure 7B).

**Figure 7:**
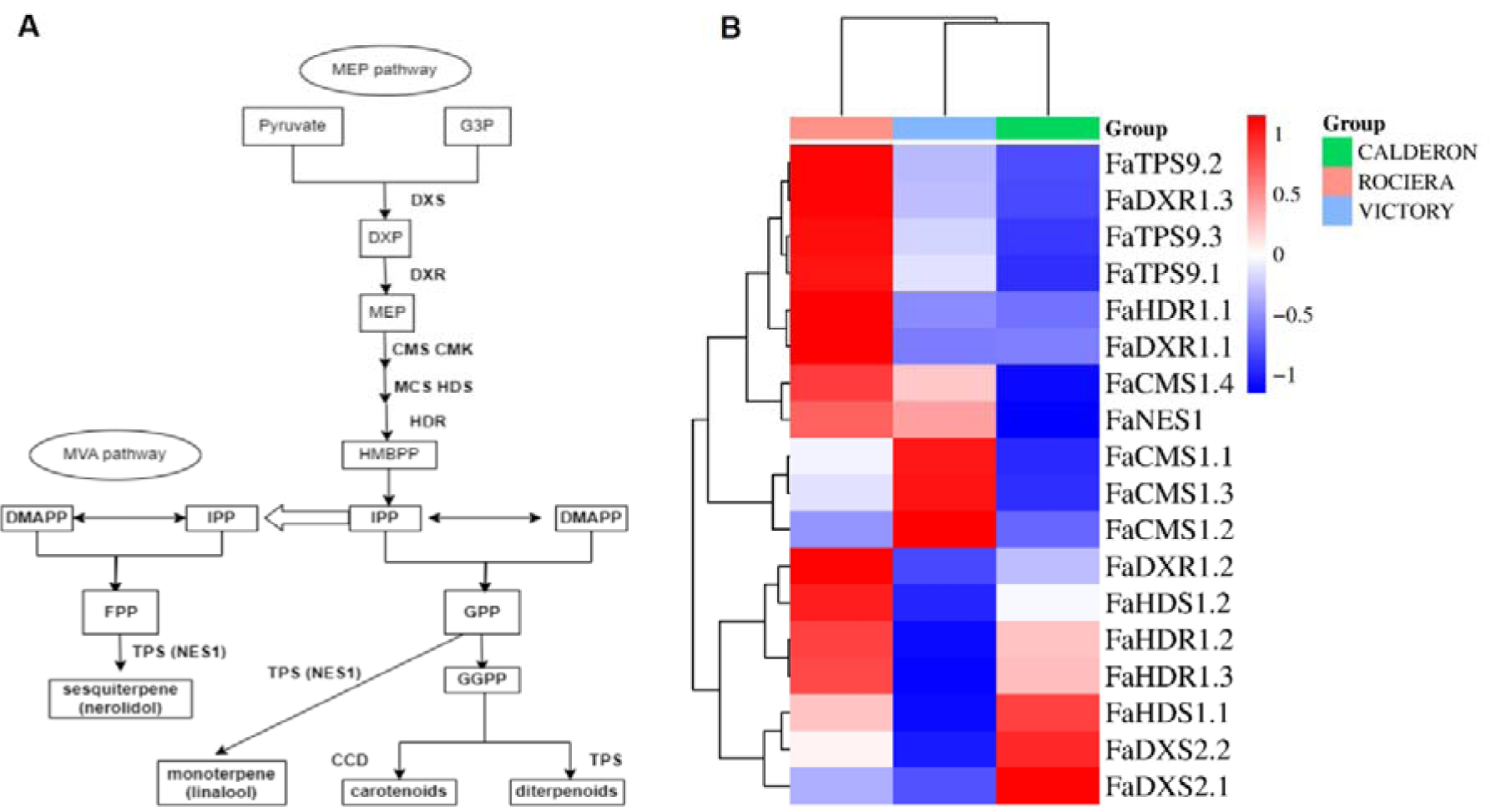
MEP pathway analysis. (**A**) Schematic diagram of the MEP pathway. Multiple enzymatic reactions are illustrated by stacked arrows. Abbreviations: DXS: 1-deoxy-D-xylulose-5-phosphate synthase; DXR: 1-deoxy-D-xylulose 5-phosphate reductoisomerase; GA-3P, glyceraldehyde 3-phosphate; CMS: 2-C-methyl-D-erythritol 4-phosphate cytidylyltransferase; CMK: 4-(cytidine 5 -diphospho)-2-C-methyl-D-erythritol kinase; MCS: 2-C-methyl-D-erythritol 2, 4-cyclodiphosphate synthase; HDS: 4-hydroxy-3-methylbut-2-en-1-yl diphosphate synthase; HDR: HMBPP reductase; TPS: terpene synthase; (**B**) Transcript level of biosynthetic genes related to MEP pathways in the fruit of ‘Rociera’ ‘Victory’ and ‘Calderon’ varieties. Darker red indicates a higher expression level, whereas darker blue indicates a lower expression level.

### Single Nucleotide Polymorphisms Linked to Differentially Expressed Genes Associated with the volatile profile in the strawberry cultivars examined

This study examined 31,677 SNPs with high impact among ‘Rociera’, ‘Calderon’ and ‘Victory’ cultivars (Table S2). Analysis identified 34 unique SNPs within genes related to volatile compound composition, affecting 14 genes with significant alterations (stop loss or stop gain alteration). In total there were 17 SNPs in ‘Rociera’ strawberry cultivar, 22 in ‘Victory’ and 24 SNPs in ‘Calderon’. Six SNPs were affecting lipoxygenase 6 protein (FxaC_14g13650, FxaC_15g14640) in ‘Victory’ and ‘Calderon’. In ‘Rociera’ cultivar, one SNP had a high impact in *FaFAD1* gene (FxaC_3g01490) with a stop lost change. There are four SNPs causing a stop loss alteration in phenylalanine ammonia-lyase 1 (FxaC_26g18320, FxaC_27g31341, FxaC_27g35250 and FxaC_28g17130) which is associated with the early step in eugenol biosynthetic pathway in ‘Victory’ and ‘Calderon’ cultivars. In ‘Victory’ and ‘Rociera’ varieties there is one high impact SNP affecting *FAEOBII* gene (FxaC_23g47390) and causing a stop loss change in emission of benzenoids II protein.

The distribution of SNPs across chromosomes and their impact on key genes such as lipoxygenase 6 and *FaFAD1* highlights the genetic variation underlying the volatile profiles of these strawberries.

## Discussion

Strawberry flavor and aroma are significantly influenced by their volatile compound content. To date, almost a thousand volatile compounds have been identified in strawberries, with esters, acids, furanones, lactones, and terpenes being the predominant and most impactful ones [49]. The volatile profile of strawberries varies widely across different cultivars [50–53]. In the biosynthesis of furanones, the *FaQR* gene is crucial for furaneol production, with *FaERF*#9 and *FaMYB98* complex enhancing its production, and *FaOMT* facilitating the creation of mesifurane. Our examination of these genes and transcription factors revealed that *FaQR* and FaMYB98 are upregulated in ‘Rociera’ compared to ‘Victory’ and ‘Calderon’, potentially leading to a higher furaneol content, as indicated by our prior research [52,53]. This is further supported by the upregulation of the *FaMYB98* transcription factor in ‘Rociera’, boosting furaneol production. *FaOMT* showed varying expression levels across the cultivars, with ‘Calderon’ showing higher expression and ‘Rociera’ a reduction. These findings offer a contrast to those of Bian *et al*. (2023), where *FaQR* and *FaOMT* exhibited similar expression patterns in the ‘Sachinoka’ variety and its somaclonal mutant [54].

Volatile esters are produced via amino acid and fatty acid pathways. The key gene *FaAAT* was found to be upregulated in ‘Rociera’, aligning with its higher ester content, such as ethyl butanoate, methyl hexanoate, and ethyl hexanoate, compared to ‘Victory’ and ‘Calderon’, leading to a more aromatic variety, as also observed by Bian *et al*. (2023) [54]. The significant role of the *FaAAT* gene in ester biosynthesis was corroborated by Zhang *et al*. (2023), who found higher *FaAAT* expression levels in the variety with increased ester percentages and odor activity values [25].

The amino acid pathway also produces phenylpropane eugenol. Genes such as *FaPAL*, *FaCOMT*, *FaCCR*, *FaCAD*, and *FaEGS2* were upregulated in ‘Rociera’, potentially leading to increased eugenol production compared to ‘Victory’ and ‘Calderon’, where these genes were downregulated. The upregulation of transcription factors FaMYB10 and FaEOBII in ‘Rociera’ supports these findings, as shown in Medina-Puche *et al*. (2015), indicating regulation of *FaCAD1* and *FaEGS2* by *FaEOBII* and *FaMYB10’s* control over *FaEOBII* expression [19]. Conversely, *FaESG1* was downregulated in ‘Rociera’, consistent with Baldi *et al*. (2018), where *FaEGS1* and *FaEGS2* exhibited opposite expression regulation [55].

Lactone biosynthesis, involving the β-oxidation of unsaturated fatty acids, showed no clear expression pattern among the cultivars, leaving lactone biosynthesis conclusions, especially regarding γ-decalactone, elusive despite significant γ-decalactone level differences found in previous studies [52,53](Leonardou *et al*., 2021; Passa *et al*., 2023). Monoterpene linalool and sesquiterpene nerolidol synthesis through the MVA and MEP pathways had *FaDXR* and *FaTPS* upregulated only in ‘Rociera’, with *FaNES1* also showing higher expression, correlating with higher linalool and nerolidol levels compared to ‘Victory’ and ‘Calderon’.

These findings shed light on the genetic variation among commercial strawberry cultivars, affecting the expression of crucial genes in volatile biosynthetic pathways and resulting in distinct volatile profiles. This study paves the way for molecular strawberry breeding aimed at improving sensory attributes. Identifying specific differentially expressed genes and SNPs linked to the biosynthesis of flavor and aroma compounds offers breeders the tools for marker-assisted selection and genomic selection to expedite the creation of cultivars that meet consumer taste preferences. Significant genes involved in pathways such as terpenoid, furanones, fatty acid, and amino acid metabolism, along with impactful SNPs on flavor-related genes, serve as key genetic markers for breeding programs focused on enhancing desirable sensory traits. Future efforts could focus on incorporating alleles that contribute to optimal flavor profiles, optimizing sweetness, acidity, and aromatic qualities.

## Acknowledgments

This research has been co-financed by European Regional Development Fund of European Union and Greek national funds through the Operational Program “Competitiveness, Entrepreneurship & Innovation” (EPAnEK), under the call RESEARCH—CREATE—INNOVATE (project code: T2EDK-01924, MIS 5129411) entitled “Development of new strawberry varieties improved in terms of aroma and taste using chemical and genetic markers—STRAWBERRY”. The authors thank Dr Eleni Liveri and Charikleia Papaioannou Msc for technical assistance in RNA extraction, Dr Anastasios Katsileros for image configuration and Professor Fotini N. Lamari for critical advice regarding the biochemical pathways.

## Supplementary Tables

Table S1. DEGs in comparisons between ‘Rociera’ vs. ‘Victory’ and ‘Rociera’ vs. ‘Calderon’

Table S2. SNPs with high impact in ‘Rociera’, ‘Victory’ and ‘Calderon’ varieties

